# Tracking and curating putative SARS-CoV-2 recombinants with RIVET

**DOI:** 10.1101/2023.02.17.529036

**Authors:** Kyle Smith, Cheng Ye, Yatish Turakhia

## Abstract

Identifying and tracking recombinant strains of SARS-CoV-2 is critical to understanding the evolution of the virus and controlling its spread. But confidently identifying SARS-CoV-2 recombinants from thousands of new genome sequences that are being shared online every day is quite challenging, causing many recombinants to be missed or suffer from weeks of delay in being formally identified while undergoing expert curation. We present RIVET – a software pipeline and visual platform that takes advantage of recent algorithmic advances in recombination inference to comprehensively and sensitively search for potential SARS-CoV-2 recombinants, and organizes the relevant information in a web interface that would help greatly accelerate the process identifying and tracking recombinants.

**Availability and Implementation:** RIVET-based web interface displaying the most updated analysis of potential SARS-CoV-2 recombinants is available at https://rivet.ucsd.edu/. RIVET’s frontend and backend code is freely available under MIT license at https://github.com/TurakhiaLab/rivet. All inputs necessary for running the RIVET’s backend workflow for SARS-CoV-2 are available through a public database maintained by UCSC (https://hgdownload.soe.ucsc.edu/goldenPath/wuhCor1/UShER_SARS-CoV-2/).

## Introduction

Recombination is a known contributor to genetic novelty in SARS-CoV-2 (Jackson *et al*., 2021; Turakhia *et al*., 2022). Recent recombinant sub-variants, such as XBB.1.5, have been identified as amongst the most transmissible variants of SARS-CoV-2 seen so far (Yue *et al*., 2023).

Identifying and tracking recombinants in a timely manner is therefore critical to help public health officials to promptly respond to emerging threats and implement appropriate control measures. Because of the epidemiological importance of recombinants, and because they violate the normal tree structure of evolution, a special naming convention (*i*.*e*., names starting with “”) is used for recombinant variants in Pango – the most widely-used nomenclature system for SARS-CoV-2 variants (Rambaut *et al*., 2020). As of February 2023, about 60 unique recombinant lineages (or about 106, if including sub-lineages) have been formally identified under this system. However, a recent large-scale study on SARS-CoV-2 recombinants found 589 uniquely identifiable recombinant lineages had emerged till May 2021 alone (Turakhia *et al*., 2022), suggesting considerable underestimation in the Pango system of the actual prevalence of detectable recombination. This is possibly because of the significant manual effort that is involved in ratifying recombinants in the present system. First, volunteers are required to collect and present evidence of recombination that they found using the pangolin-designation GitHub Issues (https://github.com/cov-lineages/pango-designation/issues). Next, another group of experts laboriously review the proposals for potential issues (such as contamination or bioinformatic errors), before formally designating recombinant lineages that meet the quality standards – a process that often takes weeks to months. In this manuscript, we present RIVET, a tool that can significantly accelerate both aforementioned steps to designate recombinant lineages. In the future, RIVET may be combined with Autolin (McBroome *et al*., 2023) to fully-automate the process of naming variants, including the special handling of recombinants. It may also help to build more accurate representations of the evolution of SARS-CoV-2 and other pathogens using phylogenetic networks.

## Results and Discussion

Figure 1A shows an overview of RIVET (Recombination Viewer and Tracker). It consists of backend and frontend components.

**Figure 1:**
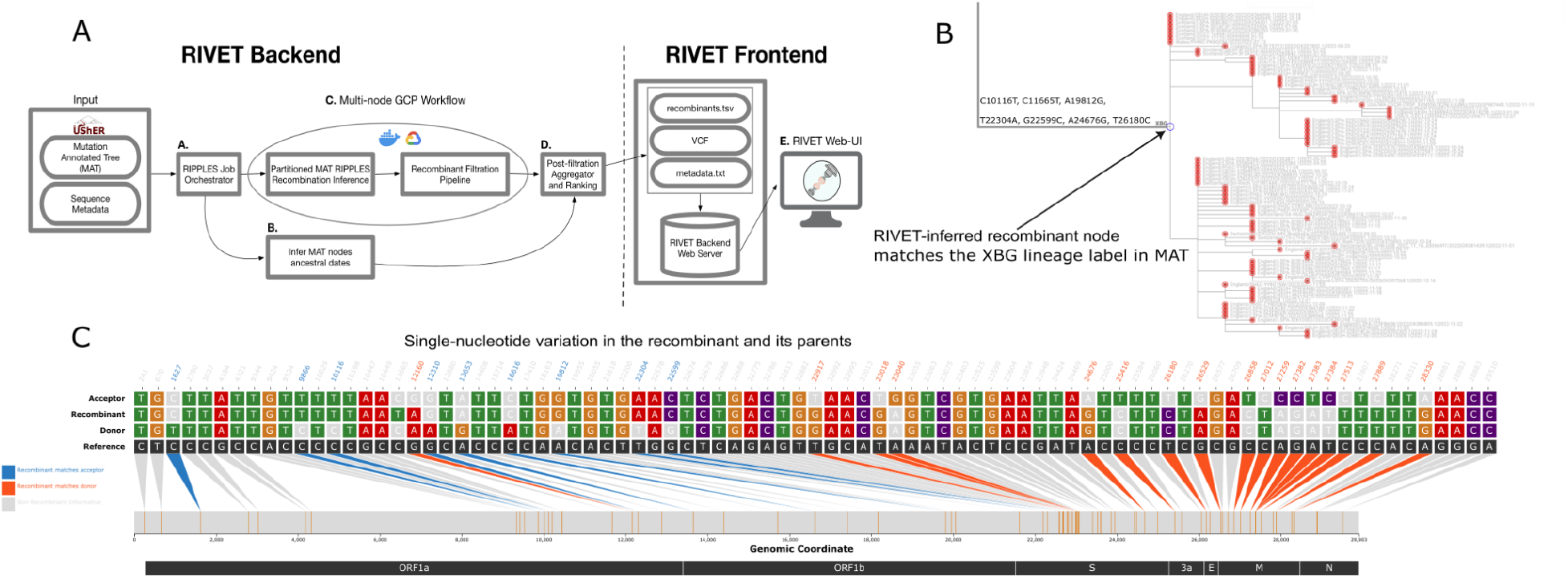
(**A**) An overview of RIVET’s backend and frontend components. (**B**) Taxonium view highlighting a recombinant node (node_1097813) identified by the RIVET workflow in the 2023-01-31 public SARS-CoV-2 MAT and (**C**) its corresponding RIVET-based SNV plot. This node in the MAT was pre-annotated as the root of the XBG lineage that was manually identified as recombinant by the Pango curation team (https://github.com/cov-lineages/pango-designation/issues/896).

RIVET’s backend (Figure 1A) consists of a workflow that starts by comprehensively searching for potential recombinants from an input mutation-annotated tree (MAT) (Turakhia *et al*., 2021) using a newly-optimized implementation of the RIPPLES software (Methods) with sensitive parameter settings (Methods). Subsequently, the workflow initiates an automated pipeline that analyzes each RIPPLES-inferred recombinant in the MAT using raw genome sequences provided to the pipeline as input for potential quality issues that may have resulted from bioinformatic, contamination or other sequencing errors. This pipeline was developed and described previously in the RIPPLES manuscript (Turakhia *et al*., 2022). The workflow also infers the dates of origin of RIPPLES-inferred recombinant nodes in the MAT using sequence metadata and Chronumental (Sanderson, 2021). This also helps to rank recombinants using an *ad hoc* growth score (Methods). RIVET’s workflow has been heavily optimized for efficient and parallel execution on the Google Cloud Platform (GCP) (Methods). For example, on a public MAT dated January 31, 2023, and consisting of 6.4 million SARS-CoV-2 sequences, the workflow produced results within an hour and with less than $3 in compute costs (Table 1). We therefore do not expect compute cost to be a major constraint to frequently update RIVET results. RIVET’s backend produces a set of output files that can be loaded to its frontend (Methods).

**Table 1:** RIVET backend runtime and cost estimates using two `n2d-highcpu-32` instances on the Google Cloud Platform (GCP) for different public MATs.

| Date of Public MAT | Number of sequences in the MAT | Number of unique recombinants discovered |  | Ripples-fast runtime | Total runtime (including QC, ranking etc.) | Estimated compute cost |
| --- | --- | --- | --- | --- | --- | --- |
|  |  | Pre-filter | Post-filter |  |  |  |
| October 31, 2022 | 6,427,951 | 1,413 | 421 | 20m 22s | 45m 25s | \$2.06 |
| November 30, 2022 | 6,497,825 | 1,452 | 441 | 20m 19s | 49m 17s | \$2.23 |
| December 31, 2022 | 6,612,971 | 1,473 | 455 | 17m 01s | 43m 10s | \$1.97 |
| January 31, 2023 | 6,716,605 | 1,470 | 453 | 23m 35s | 51m 10s | \$2.34 |

RIVET’s frontend is a Flask application (Methods) that allows for interactive analysis of backend results on a web browser, displaying a variety of useful information to aid manual curation efforts. Notably, our website (https://rivet.ucsd.edu/) that is hosted using RIVET’s frontend, displays a table of all RIPPLES-inferred trios of recombinant and parent (called donor and acceptor) nodes. The table also includes relevant information such as the inferred breakpoint intervals, parsimony score improvement resulting from the partial placement of recombinant sequences broken at breakpoints on parent nodes found by RIPPLES, current Pango lineage assignment of recombinant and parent nodes, 3SEQ-derived p-values of informative site sequence (Martin *et al*., 2010), date of origin of the recombinant, growth score, and so on. The table also displays potential quality issues for recombinants identified by the RIPPLES’ filtration pipeline. The table can be searched and sorted using any field, and the recombinant ancestors can be queried using sample identifiers. The table also links each recombinant to a Taxonium/Treenome view (Sanderson, 2022; Kramer *et al*., 2023), in which the phylogenetic contexts and genotype information of the recombinant and its parent sequences can be simultaneously visualized (Figure 1B). The frontend also provides a view of single-nucleotide variation (SNV) in the three sequences (each recombinant and its parents), with informative sites highlighted (Figure 1C). This view is inspired from *snipit* (https://github.com/aineniamh/snipit) and uses the VCF produced by the backend as input (Methods). RIVET’s frontend code also allows users to load VCFs of arbitrary sequences on a local HTTP server (instructions provided on GitHub).

RIVET analysis on public MAT dated January 31, 2023, revealed 1,470 unique recombinants, of which 453 passed all quality checks. The breakpoint distribution of high-quality recombinants which passed all automated quality checks showed increased recombination rates in the 3’ portion of the SARS-CoV-2 genome (Figure S1), consistent with the previous study (Turakhia *et al*., 2022). Intra-lineage recombination, which gets frequently overlooked, accounted for 29.8% of high-quality recombinants. RIVET inferences (such as lineages of parent sequences) of known recombinants were largely consistent with those of manual curators (Figure 1B,C). Some known recombinants (e.g., XAR) were likely missed because the public SARS-CoV-2 MAT does not include sequences from the GISAID database (McBroome *et al*., 2021). Additionally, because of minor imperfections in the heuristic to annotate root nodes of different lineages in the MAT (McBroome *et al*., 2021), we found that in some cases, the node annotated as the root of a Pango-designated recombinant lineage was 1-3 generations below the node at which RIVET inferred its corresponding recombination event (Figure S2). RIVET may also be helpful to fix these inconsistencies.

## Methods

### Optimizations and parameter settings in the RIPPLES software

The RIVET backend uses a refactored implementation of RIPPLES, called `ripples-fast`, which achieves 1–2 orders of magnitude speedup relative to the original implementation while producing identical results. The key performance optimizations include (i) amortizing computations of parsimony improvement for different breakpoint intervals, (ii) improving memory locality of the algorithm, and (iii) achieving fine-grained parallelism through vectorized instructions. (i) For every node in the mutation-annotated tree (MAT), `ripples-fast` starts by maintaining a mutation count vector that stores the number of sites along different positions in the genome at which the putative recombinant node differs from the reference. When performing partial placements, this mutation count vector is directly used to find the parsimony score improvement for each possible breakpoint interval. This eliminates the cost of traversal from root to that node in the mutation-annotated tree (MAT) to find corresponding mutations at that node. (ii) Since a single vector needs to be accessed at each node to find the parsimony score improvement, this technique of using a mutation count vector per node also improves the memory locality of the RIPPLES algorithm. (iii) Finally, because recombinants are relatively rare, `ripples-fast` utilizes the SSE-based vector instructions available on Intel Processors to test for the presence of putative recombination at multiple breakpoints in parallel. This is in addition to the multithreaded and multiprocess parallelism that was already available in RIPPLES.

By default, RIVET uses sensitive search parameters for RIPPLES (i.e, setting `--branch-length 3 --num-descendant 5 --parsimony-improvement 3`). These parameters require that a node have a branch length of at least 3 mutations and a minimum of 5 descendant tips to be considered for recombination. Additionally, the partial placement parsimony score should improve by at least 3 mutations for a node to be flagged as a potential recombinant.

### Estimating date of origin of recombinants and growth scores

Since recombinants discovered through RIPPLES correspond to internal nodes of the MAT, their origin or sampling date is not directly available through sequence metadata. However, if the sequence metadata, which contains the sampling date of each sequence in the MAT, is provided as input by the user, RIVET also launches a parallel Chronumental process (Sanderson, 2021) to build a time tree from the MAT. On RIVET’s frontend interface, users can sort the recombinant list based on the origin date to quickly review recombinants that have been inferred to have emerged recently.

Additionally, to help prioritize emerging recombinants of epidemiological interest for the purposes of recombinant lineage identification and tracking, RIVET assigns each detected recombinant a growth score and outputs a ranked list of putative recombinants. The recombinant growth metric below, G(*R*), for a recombinant node with a set of descendants *S* is defined below:

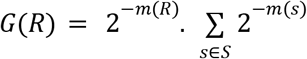

In the equation above, *m*(*R*) and *m*(*S*) correspond to the number of months (30-day intervals) elapsed since the recombinant node *R* was inferred to have originated and its descendant sequence *s* was sampled, respectively. The growth score above, *G*(*R*), is computed for each detected recombinant *R*, and the final recombinant list is ranked based on descending growth scores.

### Efficient RIVET workflow parallelization on the Google Cloud Platform (GCP) and output files

The entire RIVET backend pipeline is contained within a public Docker image that can be massively parallelized across multiple servers on Google Cloud Platform (GCP). In a YAML configuration file provided, the user can specify the number of instances and machine type to run the RIVET job. By default, we run the workflow on two `n2d-highcpu-32` instances. Upon initiating, RIVET loads the input mutation-annotated tree (MAT) and conducts a parallel search for long-branches that will be considered for the recombination search. The number of long branches is then automatically partitioned uniformly across the specified number of GCP instances. Each GCP instance searches its range of long branches in parallel for recombination events. Immediately upon completion of the search phase, an automated filtration pipeline begins on the instance to check for potential sequencing and bioinformatic quality issues with each detected recombinant. Once every GCP instance has completed both the search and filtration steps, RIVET aggregates the results from each instance locally, and ranks the recombinant results.

### RIVET’s frontend implementation details

The RIVET frontend is a Flask application (Grinberg, 2018) that loads and pre-processes the output files generated by RIVET’s backend, which includes a tab-delimited recombinant results file, a VCF file containing all the single-nucleotide variants (SNVs) of the trio sequences (recombinant, donor, acceptor) and a tab-delimited descendants file containing a mapping of all trio node ids to their respective set of descendants. RIVET utilizes *cyvcf2* (Pedersen and Quinlan, 2017), which is a Python library wrapper around *htslib* (Bonfield *et al*., 2021), to enable fast parsing of the input trio VCF file. The RIVET web interface displays the recombination results ranked by growth score in a table format where each row in the table is a detected recombinant. To see the SNVs for a particular recombinant of interest, the user can select a row to dynamically render an interactive visualization built using *d3*.*js* that displays the SNVs for the selected recombinant and its two parents, with respect to the SARS-CoV-2 reference. The plot shows all positions where at least one of the trio sequences contains a variant, however the recombinant-informative sites are highlighted where the recombinant matches the donor or the acceptor sequence, for clear visualization of the inferred breakpoint intervals. By clicking the available buttons, any view of the visualization can be downloaded in SVG format, for high-quality publication-ready figures, or copied and pasted directly into lineage proposal GitHub Issues, for example. The SNV visualization also contains several built-in interactive features, such as the ability to query and download the descendants specific to a particular node in the trio by clicking the corresponding track label.

## Acknowledgements

We thank Russell Corbett-Detig, Angie Hinrichs, Jakob McBroome, Alexander Kramer and Laura Hughes for their constructive feedback and technical support. This work was supported by the Centers for Disease Control and Prevention award BAA 200-2021-11554.

## Supplementary Figures

**Figure S1:**
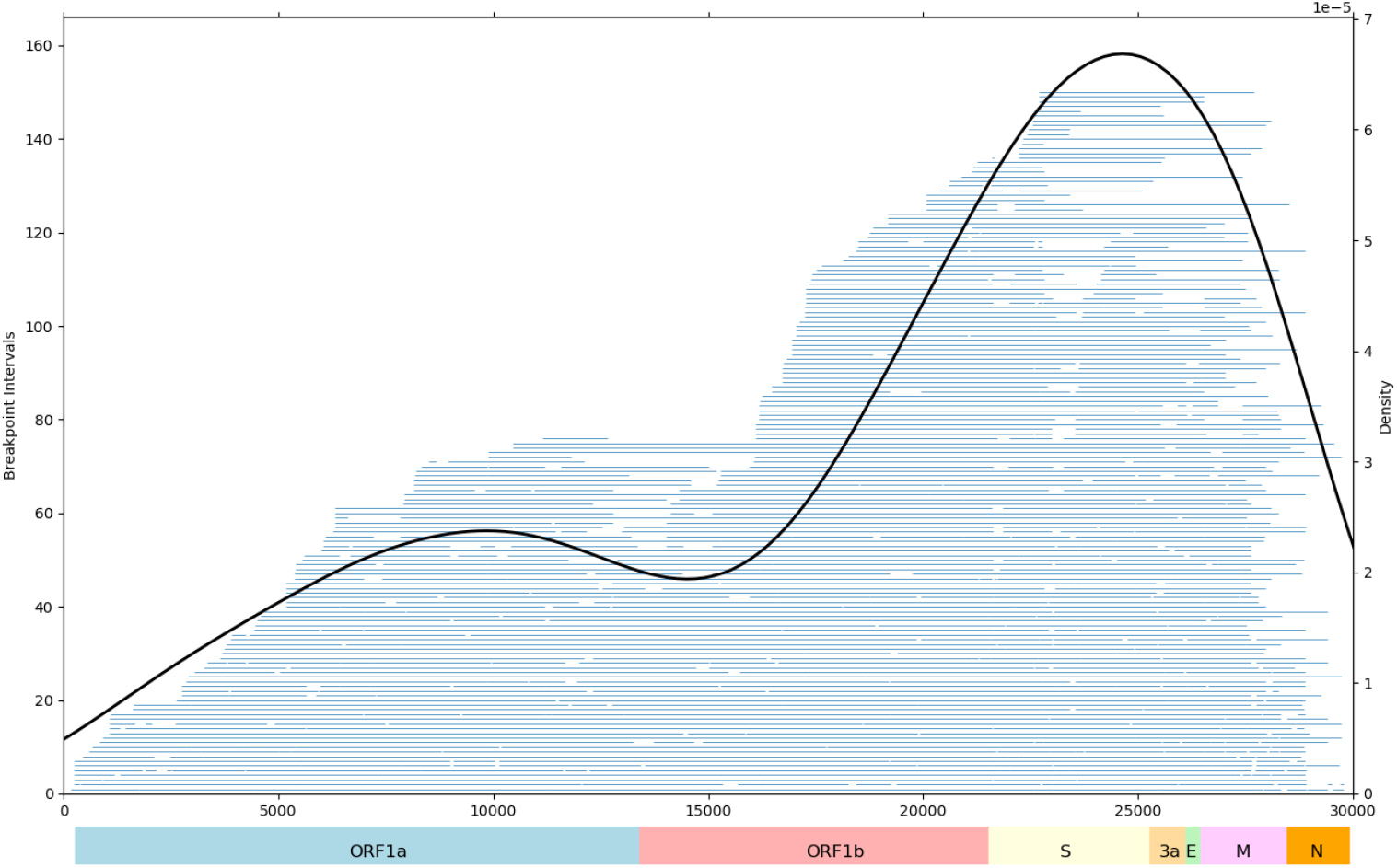
Breakpoint intervals (in blue bars) of recombinants that passed all quality filters along with a smoothed density plot of their midpoints (in black).

**Figure S2:**
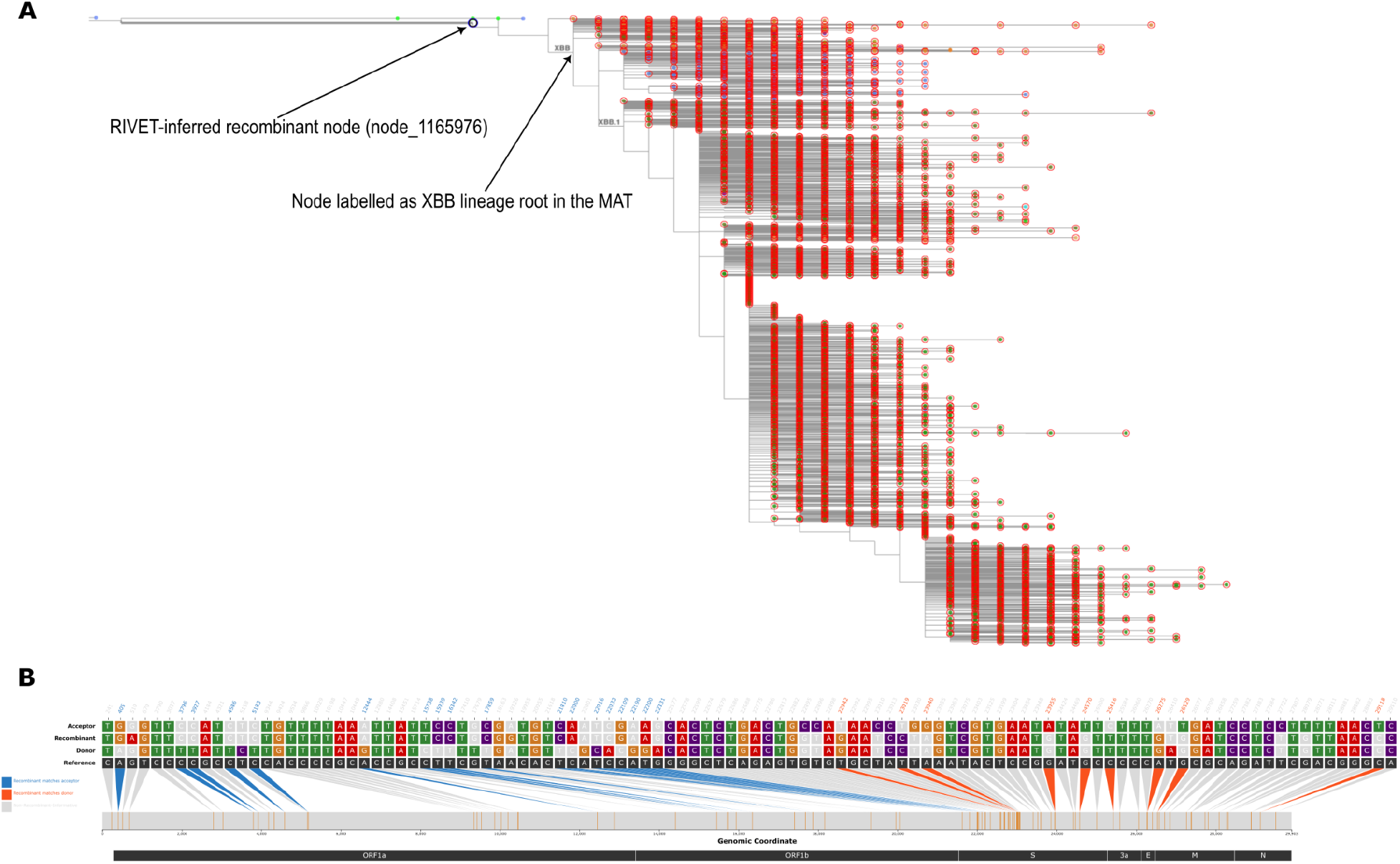
(**A**) Taxonium view highlighting the MAT node labeled node_1165976 that was inferred as a recombinant in RIVET and (**B**) its corresponding RIVET-based SNV plot. This node appears to contain the recombination event identified in the XBB lineage by the Pango curation team (https://github.com/cov-lineages/pango-designation/issues/1058). However, the root of the XBB lineage is annotated three generations below in the MAT, as evident in panel (A).

